# Using more than the oldest fossils: Dating Osmundaceae by three Bayesian clock approaches

**DOI:** 10.1101/010496

**Authors:** Guido W. Grimm, Pashalia Kapli, Benjamin Bomfleur, Stephen McLoughlin, Susanne S. Renner

**Affiliations:** Swedish Museum of Natural History, Department of Palaeobiology, Svante Arrhenius 7, Väg SE-10405 Stockholm, Sweden; Natural History Museum of Crete and Biology Department, University of Crete, P.O. Box 2208, Gr-71409, Heraklion, Crete, Greece; Systematic Botany and Mycology, University of Munich, Str. 67, 80638 Munich, Germany

**Keywords:** Bayesian inference, fossilised birth-death dating, molecular clock calibration, node dating, total evidence dating, royal ferns

## Abstract

A major concern in molecular clock dating is how to use information from the fossil record to calibrate genetic distances from DNA sequences. Here we apply three Bayesian dating methods that differ in how calibration is achieved—‘node dating’(ND) in BEAST, ‘total evidence’(TE) dating in MrBayes, and the ‘fossilised birth-death’(FBD) in FDPPDiv—to infer divergence times in the Osmundaceae or royal ferns. Osmundaceae have 13 species in four genera, two mainly in the Northern Hemisphere and two in South Africa and Australasia; they are the sister clade to the remaining leptosporangiate ferns. Their fossil record consists of at least 150 species in ∼17 genera and three extinct families. For ND, we used the five oldest fossils, while for TE and FBD dating, which do not require forcing fossils to nodes and thus can use more fossils, we included up to 36 rhizome and frond compression/impression fossils, which for TE dating were scored for 33 morphological characters. We also subsampled 10%, 25%, and 50% of the 36 fossils to assess model sensitivity. FBD-derived divergence dates were generally greater than ages inferred from ND dating; two of seven TE-derived ages agreed with FBD-obtained ages, the others were much younger or much older than ND or FBD ages. We favour the FBD-derived ages because they best match the Osmundales fossil record (including Triassic fossils not used in our study). Under the preferred model, the clade encompassing extant Osmundaceae (and many fossils) dates to the latest Palaeozoic to Early Triassic; divergences of the extant species occurred during the Neogene. Under the assumption of constant speciation and extinction rates, FBD yielded 0.0299 (0.0099–0.0549) and 0.0240 (0.0039–0.0495) for these rates, whereas neontological data yielded 0.0314 and 0.0339. However, FBD estimates of speciation and extinction are sensitive to violations in the assumption of continuous fossil sampling, therefore these estimates should be treated with caution.#

## INTRODUCTION

Calibration is the single largest problem in molecular-clock dating, influencing not only estimates of divergence times but also evaluation of the heterogeneity in substitution accumulation, since rates always derive from calibrated trees. There are many ways to calibrate the temporal significance of genetic distances. They include fossils providing minimum divergence times (Sarich and Wilson 1967, Christin et al. 2014), oceanic islands with endemic radiations providing maximum ages of cladogenesis (Schaefer et al. 2009), ancient DNA of one‘s focal group (Korber et al. 2000), host ages as maximal ages for obligate parasites (Rector et al. 2007, Bellot and Renner 2014), ratios of substitution rates between hosts and parasites (Ricklefs and Outlaw 2010), published rates from other studies (e.g., Villarreal and Renner 2014), and node ages obtained in other studies, the so-called secondary calibration approach. The most widely used of these approaches is calibration with fossils. Since the introduction of Bayesian relaxed clock approaches that implement different strict and relaxed clock models it is possible to accommodate prior notions about how tightly a fossil may fit a particular node with different prior distributions. In this framework, fossil ages can be used as point calibrations, hard minimum bounds, hard maximum bounds, soft maximum bounds, or to centre a normal distribution, a lognormal distribution, or an exponential distribution. These distributions have large effects on the obtained ages (Ho and Phillips 2009, Warnock et al. 2012), and no amount of sequence data can correct the influence of incorrect prior constraints (Yang and Rannala 2006).

The use of several fossils to calibrate nested nodes in a tree has been suggested as a possible solution (Near et al. 2005), although this does not circumvent the problem of oldest fossils having a disproportionate effect on the outcomes (Parham and Irmis 2008). A case in point is the crown age of the flowering plants (angiosperms). Numerous molecular-clock studies have constrained the relevant node to maximally 135 million years (Ma) based on a few pollen grains from Israel (dated to 132.9 Ma) that are the oldest widely accepted record of flowering plants (Brenner 1996). When left unconstrained, the angiosperm crown age is usually much older, for example, 228 (193–270) Ma (Smith et al. 2010), an estimate only slightly younger than angiosperm-like pollen from the Middle Triassic, dated to 247.2–242 Ma (Hochuli and Feist-Burkhardt 2004, 2013). This example dramatically illustrates the problems stemming from the current need to assign oldest fossils to particular nodes, in some cases a problem worsened, not alleviated, by competing ‘oldest’ fossils. Another problem with multiple fossils is that the effective calibrations may not resemble the specified calibrations because the various priors interact with each other, with the tree prior, and with other priors, such as monophyly constraints (Inoue et al. 2010, Heled and Drummond 2012).

Two Bayesian clock methods exist that do not rely on node dating via one or more ‘oldest’ fossils. They are total evidence (TE) dating (Ronquist et al. 2012a) and fossilised birth-death (FBD) dating (Heath et al. 2014). Total evidence dating uses morphological data from extant and extinct species combined with DNA data to infer node ages. Unlike node dating, total-evidence dating can be applied to a set of fossils without fixing them to specific nodes in the tree. It relies on the morphological similarity between a fossil and the reconstructed ancestors in the extant tree to infer the lengths of extinct side branches on which a fossil sits (Ronquist et al. 2012a). Total-evidence dating uses a uniform prior on the clock trees, even though trees will include terminals of different ages because of extinct side branches. The FBD approach also allows using multiple fossils, both old and young, but does not require a morphological data matrix as does TE, nor prior age densities on fossils as does ND dating. Thus far, the FBD approach has been applied to bears (Ursidae), a small clade with a fossil record from the mid-Eocene to the mid-Pliocene (Heath et al. 2014). The method uses a complex prior on the clock trees to model taxon sampling completeness and speciation, extinction, and fossilisation rates.

Here we compare node dating, total evidence dating, and fossilised birth-death in a fossil-rich and phylogenetically pivotal group of plants, the royal ferns (Osmundaceae). The royal ferns are the sister clade to all remaining leptosporangiate ferns (Pryer et al. 2004, Schuettpelz and Pryer 2007), and are incontrovertibly monophyletic. They comprise 13 species in four genera, *Osmunda*, *Osmundastrum*, *Leptopteris*, and *Todea* (Metzgar et al. 2008). The first two occur mostly in the Northern Hemisphere extending into the humid tropics, the latter two in South Africa and Australasia (Kubitzki 1990). Royal ferns have an exceptional fossil record (Miller 1971, Tidwell and Ash 1994, Bomfleur et al. 2014b, Wang et al. 2014), with more than 150 species, 17 genera, and at least three extinct families ranging from the Permian to Neogene. The fossils are mostly foliar remains, but many are anatomically preserved (permineralised) axes.

A molecular clock model for ferns that included two Osmundaceae and was calibrated using numerous fossils, inferred a stem age for the Osmundaceae of 320 Ma (Late Carboniferous; Pryer et al. 2004). This result, however, was obtained with the crown age of (the two) Osmundaceae constrained to 206 Ma (Late Triassic) based on a point calibration derived from the rhizome fossil *Ashicaulis herbstii* from the Late Triassic of Argentina (Tidwell and Ash 1994). This fossil is a doubtful member of modern (crown) Osmundaceae (Miller 1971, Tidwell and Ash 1994, Wang et al. 2014). Within-Osmundaceae divergence events have not been the focus of a previous molecular clock study. The discovery of an unusually well preserved 182–190 million year-old *Osmunda* fossil from Sweden (Bomfleur et al. 2014a) recently prompted a critical re-evaluation of the rhizome fossils of modern Osmundaceae (Bomfleur et al. 2014b). This reappraisal (involving morphological data matrices) provided the opportunity to estimate the timing of major evolutionary events within the family, using both TE and FBD molecular clock dating. These two methods are the first to fully integrate fossils and molecular data, modelled as representing a single macro-evolutionary process. They may result in older divergence times than traditional node dating (Ronquist et al. 2012a: Hymenoptera, Heath et al. 2014: Ursidae) and can identify likely erroneous calibration fossils, as was the case for four of seven Hymenoptera calibration points (Ronquist et al. 2012a). To explore this possibility, we also applied traditional node dating, using five oldest fossils alone or in combination.

Our phylogenies are based on ∼8.6 kb of plastid DNA data for 13 extant species, 33 morphological characters scored for 19 permineralised rhizomes (data from Bomfleur et al. 2014b), and 17 frond compression/ impression fossils, assessed on the basis of key autapomorphic features for the present study. Incorporating fossils into tree models as undertaken here in principle allows one to infer speciation and extinction rates with reasonable confidence as shown with simulated data by Heath et al. (2014) and is probably an improvement over inferring these rates only from neontological data. To test how rates inferred from the FBD process would differ from those obtained with just the 13-species tree of living Osmundaceae, we carried out a TREEPAR analysis (Stadler 2011).

## MATERIALS AND METHODS

### DNA Sampling, Alignment, and Phylogenetic Analyses

We relied on the plastid DNA matrix of Metzgar et al. (2008), which consists of the protein-coding *rbc*L, *atp*A, and *rps*4 genes and the spacers *rbc*L*-acc*D*, atp*B*-rbc*L*, rps*4*-trn*S, *trn*G*-trn*R, and *trn*L*-trn*F. A few regions were excluded due to ambiguities in the alignment (29 bp in *rbc*L*-acc*D, 129 bp in *rbc*L*-atp*B, 51 bp in *rps*4*-trn*S, 39 bp in *trn*G*-trn*R, and 146 bp in *trn*L*-trn*F) and missing data (*rps*4), resulting in a matrix of 13 species and 8616 nucleotide positions. We excluded the outgroup, Gleicheniales, because of long-branch attraction (Bomfleur et al. 2014b) and follow these authors in rooting Osmundaceae between *Todea/Leptopteris* and *Osmunda/Osmundastrum* instead of between *Osmundastrum* and the remaining three genera (Yatabe et al. 2005, Metzgar et al. 2008). Sampling covers the seven species of *Osmunda* from the three subgenera *Claytosmunda*, *Osmunda*, and *Plenasium*; the single species of *Osmundastrum*, three of the ten species of *Leptopteris*, and both species of *Todea* (the second species lacked morphological data and was therefore excluded from some analyses). Osmundaceae generic and subgeneric classification schemes (Miller 1971, Yatabe et al. 2005) are shown in Fig. S1 in the online supplement archive (OSM; see *Documentation*). Species names and authors, herbarium vouchers, deposition in herbaria, and geographic origin of material are provided by Metzgar et al. (2008).

Phylogenetic analyses relied on maximum likelihood (ML) as implemented in RAXML 8.0.22 (Stamatakis 2014), using the GTR + Γ substitution model. We ran un-partitioned analyses as well as partitioned ones in which RAxML found separate substitution rates for the two genes (*rbc*L and *atp*A) and the five spacers. Support for the ML topology was assessed with the rapid bootstrap/automatic bootstop implementation in RAXML (Stamatakis et al. 2008, Pattengale et al. 2009).

### Node Dating with Oldest Fossils, Total Evidence Dating, and Fossilised Birth-Death Dating

The 36 fossils included in our study (listed in Table S1 in OSA, along with supporting references, localities, ages and other details) are unambiguous members of modern Osmundaceae. The 19 rhizomes of Jurassic to Neogene age were included in a morphological matrix by Bomfleur et al. (2014b). The 17 Triassic to Neogene fronds, including 14 *Osmunda* spp., two *Osmundopsis* spp. and *Todea amissa*, are placed within modern Osmundaceae based on lineage-diagnostic fertile or sterile features (references in Tables S1 and S2 in OSA). The species-rich fossil genera *Cladophlebis* (including >80 described fossil species) and *Todites* (including >35 fossil species), usually identified as osmundaceous foliage fossils (Tidwell and Ash 1994), possess insufficient diagnostic characters for unambiguous assignment; some may represent one of the numerous extinct lineages of Osmundales.

The oldest fronds (*O. claytoniites*) with features diagnostic of modern members of *Osmunda/Osmundastrum* are of Late Triassic age and were associated with the root node (Node A in Fig. 1). Jurassic fronds were also associated with the root because they exhibit only symplesiomorphies of the *Osmunda/Osmundastrum* clade or parallelisms limited to its subclades. Fronds with apomorphic characters diagnostic of *Osmunda* or its subgenera were associated with Node C and D (Fig. 1). They are of Early Cretaceous and younger ages. *Todea amissa* (early Eocene, Argentina) has fronds characteristic of *Todea* and was thus linked to Node B.

For node dating in BEAST V1.8 (Drummond et al. 2012), we used the five oldest fossils as constraints. We ran BEAST with the tree topology fixed to the ML topology, a partitioning scheme chosen by JMODELTEST (Darriba et al. 2012) comparing 24 models under the Bayesian information criterion: *atp*B*-rbc*L HKY+I, *atp*A 1^st^ codon HKY+I, *atp*A 2^nd^ codon HKY+I, *atp*A 3^rd^ codon HKY, *rbc*L*-acc*D HKY+I, *rbc*L1 1^st^ codon HKY, *rbc*L 2^nd^ codon HKY, *rbc*L 3^rd^ codon HKY + I, *rps*4*-trn*S HKY+Γ, *trn*G*-trn*R HKY+Γ, *trn*L*-trn*F HKY+Γ, an uncorrelated lognormal clock model, and an exponential prior age distribution on each constraint, with the offset value set to the minimum age of the stratum comprising the respective fossil and the standard deviation so that the maximum age of the stratum was included in the 97.5% quantile. One frond fossil was assigned a point age of 52 Ma based on Carvalho et al. (2013; Table S1). Each analysis was run twice for 2*10^7^ generations, sampling trees every 1000^th^ generation. The effect of the priors on the posterior values was evaluated by running an additional analysis without the data (the DNA alignment). The parameters of all runs were evaluated using TRACER v. 1.5 (Rambaut and Drummond 2009) to confirm that (i) each Markov chain reached stationarity, (ii) the effective sample sizes (ESS) were >200 for all optimised parameters, and (iii) independent runs produced convergent results. In each Markov chain Monte Carlo (MCMC) run, the first 10% of the samples were discarded as burn-in; the remaining samples were summarised in TREEANNOTATOR (part of the BEAST package) and visualised using FIGTREE (Rambaut 2014).

For total-evidence dating in MRBAYES 3.2 (Ronquist et al. 2012b) we used the same DNA matrix as before (except that *Todea papuana* was excluded because it lacked too many rhizome characters were unavailable from the literature) plus a morphological matrix with 33 characters for the 19 rhizome fossils (Bomfleur et al. 2014b). We used two data partitions, with GTR+Γ for the DNA matrix and Mk model for the morphological matrix (Lewis 2001), as implemented in MRBAYES. No topological constraints were employed, and the tree prior was uniform (Ronquist et al. 2012a). Two rhizomes were assigned a point age of 16 Ma based on Pigg and Rothwell (2001; details shown in Table S1); all other fossils were assigned uniform distributions. Three independent MCMC runs, with eight heated chains each (temperature parameter set to 0.1), were run for 2*10^7^ generations, sampling trees every 2000^th^ generation. Convergence was checked as for the BEAST run, and in each MCMC run, the first two million generations of the samples were discarded as burn-in and the remainder summarised and visualised.

For FBD dating, we relied on FDPPDIV V.3f18ed7db29f985f12785b948998f42dfa8323af (available at https://github.com/trayc7/FDPPDIV), with the ML topology as input tree (Fig. S1). FDPPDIV treats the topology and branch lengths of the extant species tree as given and does not permit assignment of separate substitution rates to specific data partitions. As required for FDPPDIV, the age of each fossil was drawn prior to analysis from a uniform distribution of its age range (ranges are given in Table S1). We performed two MCMC runs of 2*10^7^ MCMC generations, sampling every 1000^th^ generation. Log files were then compared in Tracer to check that convergence had been reached.

**Figure 1.**
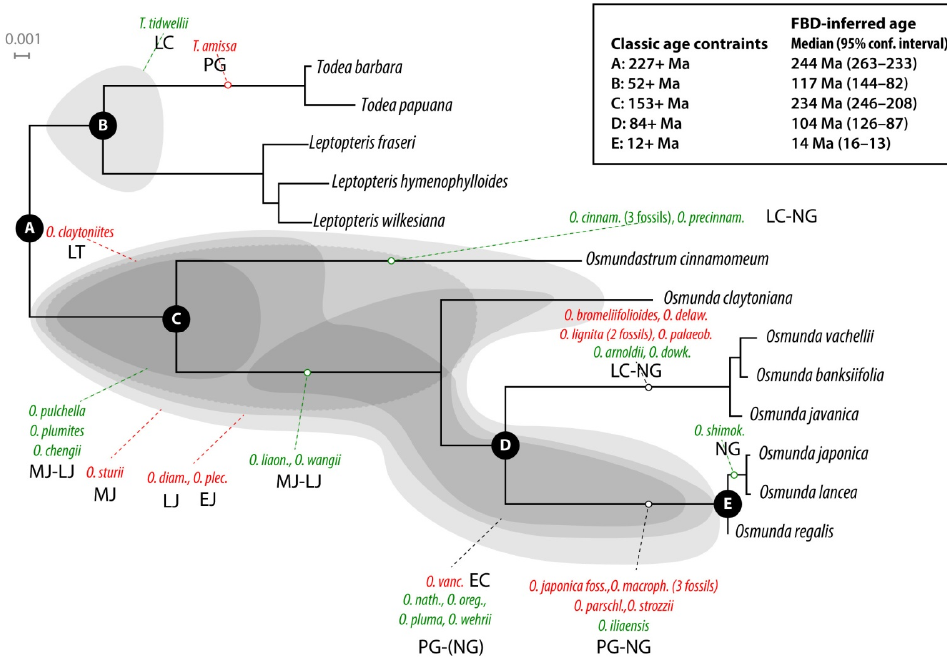
Fossil record of modern Osmundaceae mapped on a maximum likelihood tree from a plastid DNA matrix of 8616 aligned positions analysed under the GTR + Γ model with two data partitions. Rhizomes in green, fronds in red, with fossils either assigned to a branch (stippled lines) or a tree portion (shaded areas). Nodes (A–E) for which minimum age constraints were used are shown in the inset. Stratigraphic ranges of fossils abbreviated as follows: LT = Late Triassic; EJ, MJ, LJ = Early, Middle, and Late Jurassic; LC = Late Cretaceous; PG = Paleogene; NG = Neogene.

### Effects of Using Rhizomes vs. Fronds and Different Fossil Percentages

To be scored for the morphological matrix required for TE dating, fossils need sufficient characters. The frond and rhizome fossils used here differ greatly in this respect, with rhizome fossils having many more codable characters. This meant that for TE dating only rhizome fossils could be used. To be able to compare TE results with FBD results, we carried out additional FBD runs that used only the 19 rhizomes or only the 17 fronds. We also ran FBD analyses for which we randomly drew 10% (4 fossils), 25% (9 fossils), or 50% (18 fossils) of the 36 fossils, with each drawing repeated 10x. Otherwise, these runs relied on the same settings as used for the full data sets.

### Inferring Speciation and Extinction with TREEPAR vs. the FBD Approach

To quantify diversification through time we estimated tree-wide speciation and extinction rates using the R package TREEPAR (Stadler 2011) on just the extant species tree and the FBD process on the tree with all 36 fossils. FBD assumes a constant rate model, so we fitted only this model of the many available in TREEPAR.

### Documentation

Input and output files (used matrices, trees, calibration results), supplement figures and tables are included in an online supplement archive (OSA) hosted at www.palaeogrimm.org/data/Grm14_OS.zip.

## RESULTS

### Dating with only Oldest Fossils, Total Evidence, and the Fossilised Birth-Death Method

Figure. 1 schematically shows the placement of the 36 fossils used in the FBD approach (which include the 19 used in TE dating) on an ML tree from the partitioned 8616-nucleotide matrix for the 13 extant species. It also shows the five nodes (A–E) for which minimum age constraints were used in the node dating approach (in BEAST). All branches received 100% bootstrap support (Fig. S1 in OSA). Results from node dating with oldest fossils (using partitioned or unpartitioned DNA data) are shown in Table 1. The average median substitution rate (over all branches) was 1.79*10^-4^ in the unpartitioned analysis and 1.48*10^-4^ with two partitions, close to the 1.33*10^-4^ optimised under the FBD model (Table 1). Regardless of data partitioning, the ages of nodes inferred via node dating are younger than those inferred from FBD (Table 1; Table S3 in OSA), with differences of up to 82 Ma (in the case of the *Osmunda* crown age) or 63 Ma (for the *Todea/Leptopteris* divergence).

**Table 1.** Divergence ages (median values and 95% highest posterior density intervals) and optimised substitution rates obtained with fossilised birth-death dating (FBD), total evidence (TE) dating, or node dating (ND) with just the five oldest fossils as minimum age constraints (nodes A–E in see Fig. 1). The DNA matrix, substitution model and partitioning schemes are described in *Materials and Methods*. FDPPDIV does not allow data partitioning. The chronogram in Fig. 2 visualises the FBD results; the TE chronogram is shown in Fig. S2 in OSA; node dating chronograms not shown. Similar ages from the three methods are indicated in bold.

| Node | FBD | TE | ND<br>unpartitioned | ND<br>partitioned |
| --- | --- | --- | --- | --- |
| Root node | 243 (264–233) | 185 (187–183) | 229 (234–227) | 228 (231–227) |
| *Todea/Leptopteris* split | **116 (132–100)** | **113 (150–66)** | 53 (55–52) | 55 (62–52) |
| *Osmunda/Osmundastrum* split | 238 (264–233) | 182 (186–176)<sup>a</sup> | 156 (164–153) | 176 (201–153) |
| *Osmunda* crown | **133 (150–119)** | **143 (171–110)<sup>b</sup>** | 107 (129–91) | 104 (115–95) |
| Subgenera *Plenasium/Osmunda* split | 111 (126–98) | [see above]<sup>b</sup> | 84 (86–84) | 84 (86–84) |
| *Todea* crown | **13 (23–4)** | ~ | **12 (27–2)** | 9 (20–3) |
| Subgenus *Leptopteris* crown | **27 (34–21)** | 49 (105–9) | **25 (38–12)** | 19 (24–14) |
| Subgenus *Plenasium* crown | **9 (13–5)** | 84 (123–62) | 12 (27–4) | **7 (10–4)** |
| Subgenus *Osmunda* crown | **13 (15–13)** | 21 (58–2) | **12 (13–12)** | **12 (13–12)** |
| Substitution rate | 1.33*10<sup>-4</sup><br>(1.19–1.46) | 1.59*10<sup>-4</sup>(<sup>c</sup>) | 1.79*10<sup>-4</sup>(<sup>c</sup>) | 1.48*10<sup>-4</sup>(<sup>c</sup>) |

<sup>a</sup> Age for the MRCA of *Osmunda* and *Osmundastrum* = MRCA of all extant Osmundaceae because of suspected misplacement in TE run (Fig. S2).
<sup>b</sup> MRCA of subgenera *Osmunda* and *Plenasium* = MRCA of all three subgenera of *Osmunda* (inter-subgeneric relationships not resolved in the TE consensus tree)
<sup>c</sup> Averaged over all branches, and partitions.

Results from FDPPDIV (FBD) dating with either the 19 fossil rhizomes or the 17 fronds differed little (Table S3), and here we concentrate on results employing all 36 fossils (Fig. 2). With 36 fossils, the FBD method indicated an Anisian (Middle Triassic) age of 243 (264–233 95% HPD) Ma for the Osmundaceae crown group (Table 1), with subsequent radiations dated as 238 (262–226; Ladinian, Middle Triassic) Ma (*Osmunda* ↔ *Osmundastrum*) and 116 (132–100; Early Cretaceous) Ma (*Leptopteris* ↔ *Todea*). Divergences within *Osmunda* are placed in the late Mesozoic at 133 (150–119) and 111 (126–98) Ma (earliest to mid-Cretaceous). FBD runs that used 10%, 25%, or 50% of the total fossils, showed that 50% were sufficient to obtain results similar to those recovered with all 36 fossils (Table 2; see Table S4 in OSA for detailed results). With four or eight fossils, divergence ages (on average) became 20–70% younger than when using all 36 fossils. However, divergence times from the subsampled fossils are not all younger than those obtained with the full set of fossil constraints.

**Table 2.**
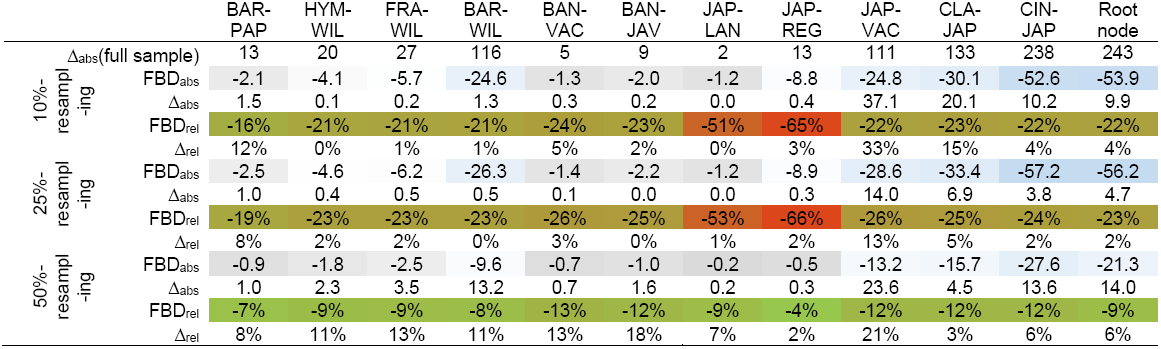
Stability of FBD dating inferences using subsets of 4, 9, or 18 fossils, drawn randomly from the total set of fossils (10%, 25%, 50% resampling), with each subsampling repeated 10x. Shown are the absolute (FBD_abs_) – grey: older, blue: younger – and relative differences (FBD_rel_) – lush green: no difference, red: strong difference – to the divergence ages as inferred using all 36 fossils and the FBD approach (Fig. 2), and, without colour gradients, the difference between both approaches (Δ_abs_; Δ_rel_). Δ_abs_ for the complete fossil data set is shown for comparison in the first row. Nodes are labelled by taxon-pairs, species abbreviated by three letters and refer to the following MRCA: BAR-PAP = MRCA of *Todea*; HYM-WIL = MRCA of *Leptopteris hymenophylloides-L. wilkesiana*; FRA-WIL = MRCA of *Leptopteris*; BAR-WIL = MRCA of *Leptopteris-Todea*; BAN-VAC = MRCA of *Osmunda banksiifolia-O. vachellii*; BAN-JAV = MRCA of subgenus *Plenasium*; JAP-LAN = MRCA of *Osmunda japonica-O. javanica*; JAP-REG = MRCA of subgenus *Osmunda*; JAP-VAC = MRCA of subgenus *Osmunda*-*Plenasium*; CLA-JAP = MRCA of *Osmunda* s.str. (subgenera *Claytosmunda, Osmunda*, and *Plenasium*); CIN-JAP = MRCA of *Osmunda* s.l. (incl. *Osmundastrum*)

The TE runs with the 19 rhizome fossils added to the DNA matrix yielded a majority rule consensus tree (Fig. S2 in OSA) with seven nodes also found in the DNA tree (Figs 1 and S1); the ages of which could thus be compared between methods (Table 1). Two of the seven TE-derived ages agreed with the FBD-obtained ages, the others were much younger or much older than ND or FBD ages. Some deep nodes came out younger because the oldest rhizome fossil is 40 Ma younger than the oldest frond fossil Fig. S2 shows both the TE-derived chronogram and an FBD-derived chronogram with just rhizome fossils). Jurassic fossils, including *Osmunda pulchella*, which has characters intermediate between *Osmundastrum* and *Osmunda*, were placed basal to clades comprising the extant species of these genera, matching the fossils’ plesiomorphic or ambiguous morphological characters, and this resulted in a Jurassic root age (Table 2). Fossils with more apomorphic features than their modern relatives, such as the Paleogene fossils of subgenus *Plenasium*, resulted in very old crown ages for this subgenus. The *Osmundastrum* lineage was resolved as sister to the *Todea/Leptopteris* lineage, whereas in the DNA-only tree it is sister to *Osmunda*.

The geographic history of Osmundaceae through time is visualised in Fig. 2 next to the FBD time tree. The *Osmunda/Osmundastrum* lineage was established in the Southern Hemisphere (Antarctica) by the Late Triassic, but the major radiations appear to have taken place in the late Mesozoic and Paleogene of the Northern Hemisphere. The *Todea/Leptopteris* lineage, today confined to southern Africa (*Todea*) and Australasia (both genera), is represented in the fossil record by one Northern Hemisphere Early Cretaceous rhizome and one frond from the Paleogene of South America.

### Speciation and Extinction Rates from Neontological Data vs. with Fossils Included

With the FBD method, the speciation rate (λ) was estimated as 0.0299 (0.0099–0.0549), the extinction rate (μ) as 0.0240 (0.0039–0.0495), indicating a high turnover and relatively slow diversification rate (0.006 per million years). The fossil recovery rate (ψ), which models how many lineages (extinct and extant) are covered by the fossil sample, was ψ = 0.01531, meaning that there is a 31% probability that a species will be represented in the fossil record. Inference (using TREEPAR) of speciation and extinction rates from just neontological data gave slightly higher speciation (λ = 0.0314) and extinction rates (μ = 0.0339; Table 2), but both values fell within the confidence intervals of the FBD-inferred rates.

**Figure 2.**
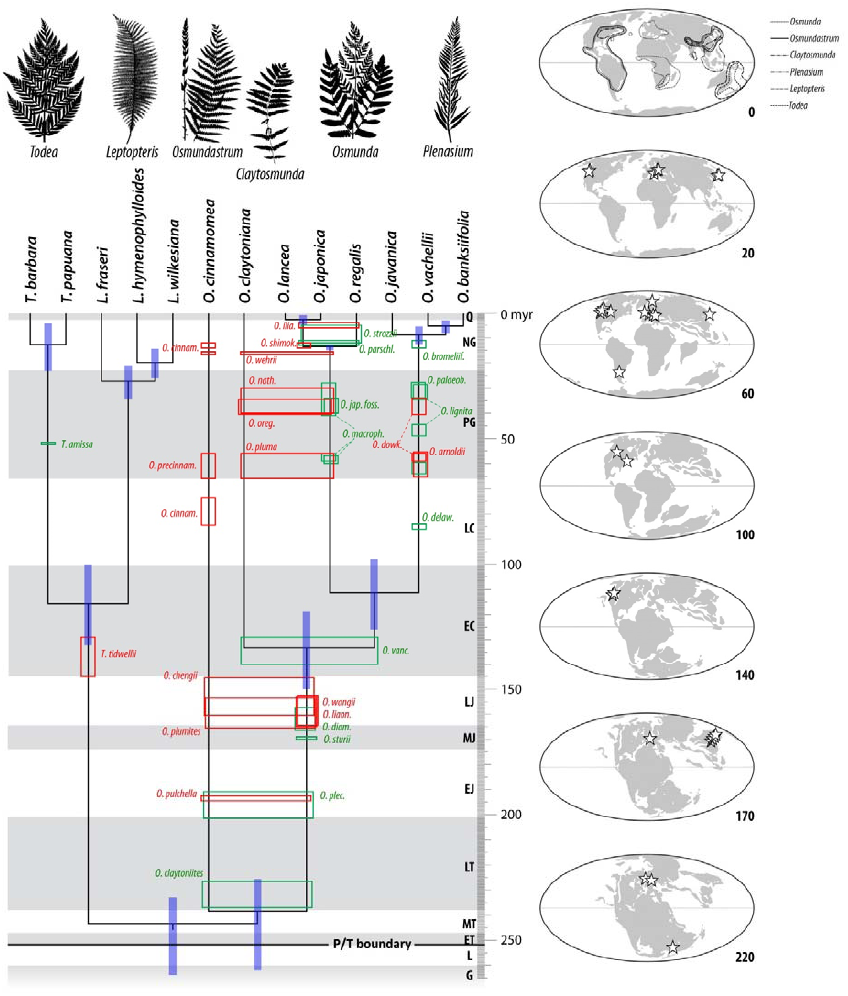
Chronogram for Osmundaceae as inferred from the fossilised birth-death approach, with the inferred placement of the 36 fossils used (red = fronds; green = rhizomes). Blue bars represent 95% highest posterior density intervals. The palaeogeographic locations of the fossils are shown on the maps to the right. Geological periods and epochs abbreviated: G = Guadalupian, L = Lopingian (Permian); ET, MT, LT = Early, Middle, and Late Triassic; EJ, MJ, LJ = Early, Middle, and Late Jurassic; EC, LC = Early and Late Cretaceous; PG = Paleogene; NG = Neogene; Q = Quaternary.

## DISCUSSION

### Comparison of the Three Bayesian Calibration Approaches

The first method to implement a full integration between fossil and molecular data was the total evidence (TE) approach of Ronquist et al. (2012a). In their original study, Ronquist et al. included even poorly preserved fossils (14 of their 45 fossils could only be coded for 4–10% of the 343 morphological characters), yet recovered topologies did not change and the median node ages were similar to those obtained with subsets excluding these poorly preserved fossils. In our study, the 19 rhizome fossils used in the TE analyses, coded for up to 33 morphological traits (Bomfleur et al. 2014b), were placed as expected based on the analyses by Bomfleur et al. We have no morphological matrix for the frond fossils, so they could not be used in TE dating. The strengths of TE dating, the parallel optimisation of node heights and phylogenetic placement of fossils are also its major weaknesses: The method forces workers to carefully analyse and score morphological traits, but where comprehensive morphological data are unavailable, it cannot be used.

Traditional Bayesian node-dating-using-oldest-fossils (Drummond et al. 2006) does not require a morphological matrix and has proven its power in hundreds of studies. Where possible, workers have used several fossils to calibrate nested nodes in a tree, attempting to reduce disproportionate influences of single fossils (Parham and Irmis 2008). Recent work, however, shows that the *ad hoc* priors placed on the ages of fossils in traditional Bayesian calibration (e.g., in the software BEAST) can be problematic because the probability distribution for the age of each calibrated node comes from *both* the node-specific calibration prior and the tree-wide prior on node ages. This leads to incoherence in the model of branching times on the tree (Heled and Drummond 2012, Warnock et al. 2012, Heath et al. 2014, Silvestro et al. 2014). This problem is avoided by applying a birth-death process to un-calibrated nodes conditioned on the calibrated nodes (Yang and Rannala 2006), which seems a more realistic representation of the lineage-diversification process and is achieved in the FBD approach (Heath et al. 2014).

The FBD approach does not require a morphological matrix and can, in principle, use the entire fossil record of a focal group, which greatly reduces the impact of unrepresentative (or misinterpreted) oldest fossils. Not having to compile a morphological matrix will be welcomed by phylogeneticists interested in divergence times, yet without expertise and resources for morphological work on living and fossil taxa. For many groups, perhaps especially plants, building morphological matrices including fossil and extant taxa may not be feasible (as was the case here for Osmundaceae fronds). A weakness of the FBD approach might be that it does not permit assignment of separate substitution rates to separate data partitions. Partitioning of substitution models, however, can cause statistical problems in clock dating (dos Reis et al. 2014). In our data set, there were no strong differences between ages inferred from un-partitioned BEAST clock dating runs and partitioned ones (Table 1).

### Implications of the Inferred Divergence Scenario for the Evolution of Osmundaceae

Initial radiations in the ferns took place in the mid-Palaeozoic (Taylor et al. 2009). Monilophytes, the group incorporating whisk ferns (Psilotales), adder‘s tongue ferns (Ophioglossales), horsetails (Equisetales), marattioid ferns (Marattiales), and leptosporangiate ferns (Polypodiales), have a fossil record extending back to at least the Late Devonian (Taylor et al. 2009). Osmundaceae in the broad sense (including the primitive Thamnopteroideae) were established in the Permian, with many well-preserved (permineralised) rhizomes from the Middle to Late Permian of Australia and Russia (Gould 1970, McLoughlin 1992) and members of their sister family Guaireaceae recorded from coeval strata in South America (Herbst 1981). The Guaireaceae and Osmundaceae emerged during a phase of stepwise global warming in the wake of the Late Palaeozoic Ice Age, with Thamnopteroideae then becoming extinct during the end-Permian biotic crisis. The core Osmundaceae persisted in moist temperate to tropical climates to the present and extended to high latitudes during phases of greenhouse climates in the Mesozoic and Paleogene (Collinson 2002).

Given this phylogenetic and fossil background, we contend that the older Osmundaceae divergence times inferred with the FBD method (and also partly the TE approach; Table 1) are more consistent with the palaeontological record than the mostly younger ages inferred with node-dating-using-oldest-fossils. Osmundaceous fronds (of poorly understood affinity; Table S2: *Todites*, *Rooitodites*, *Birtodites* and *Elantodites*) are common from the Late Triassic onwards. The FBD-inferred Middle to Late Triassic stem ages of the lineages *Osmunda/Osmundastrum* and *Leptopteris/Todea* are consistent with these palaeontological dates; these lineages, apparently beginning to diverge in the aftermath of the end-Permian mass extinction (Tidwell and Ash 1994, Taylor et al. 2009; Fig. 2). The preferred chronogram (Fig. 2) further indicates segregation of *Todea* and *Leptopteris* and of the three subgenera of *Osmunda* during the mid-Cretaceous and radiation and establishment of extant species in the Neogene. Modern Osmundaceae appear to have originated in the humid temperate belt of southern Gondwana (see maps in Fig. 2). The modern distribution of the single species of *Osmundastrum* (= *Osmunda cinnamomea*) includes humid climate tracts of South America, eastern North America and East Asia. Fossil evidence places *Osmundastrum* in Canada by the Late Cretaceous (Serbet and Rothwell 1999), so range expansion to both hemispheres may have occurred during the more humid phases of the mid-Mesozoic.

An implication of the inferred Late Triassic crown age of *Osmunda/Osmundastrum* is that Early to Middle Jurassic rhizomes, which are intermediate between *Osmundastrum* and *Osmunda*, represent stem group taxa of either *Osmundastrum* or *Osmunda*. For the recently described 182–190 million year-old *Osmunda pulchella* from Sweden (Bomfleur et al. 2014b), the divergence times obtained here suggest that it may represent an early precursor of the *Osmundastrum* lineage. The ‘*Osmundastrum* precursor’hypothesis is one of three alternatives that can be inferred from the set of analyses conducted by Bomfleur et al. (2014b). Our molecular (FBD) dating also provides a time frame for the morphological innovations in modern Osmundaceae. The post-Cretaceous increase in shaded, moist niches in angiosperm-dominated forests may have provided improved ecological conditions for Osmundaceae, particularly for a lineage such as *Leptopteris*, which cannot survive without permanent moisture (Brownsey and Perrie 2012).

### Inferring Speciation and Extinction Rates with the FBD Approach

Incorporating many fossils allows inferring speciation and extinction rates with confidence as shown with simulated data by Heath et al. (2014). In our Osmundaceae data, speciation and extinction rates from the FBD approach and the neontological data (with TREEPAR) were similar (Table 3), with the confidence intervals around the FBD rate bracketing the TREEPAR rates. With very few tip species, TREEPAR sometimes infers higher extinction than speciation rates, which seems to have been the case here (13 tip species). A parameter that may influence (and distort) the inferred speciation and extinction rates is the sampling rate of the living species, which is set to ‘1’in FDPPDIV. Future within-species sampling may reveal that some *Osmunda* forms include multiple species. For instance, the extremely widespread *O. (Osmundastrum) cinnamomea* (∼5 common synonyms) and *O. regalis* (∼10 synonyms) show intra-specific morphological variation (not covered in our molecular data) that is comparable to inter-specific diversity in other Osmundaceae. This would mean that species sampling in this study might not have been complete. Estimates of speciation and extinction are also sensitive to violations in the assumption of continuous fossil sampling (T. Heath, pers. comm., Sep. 2014), and the fossil sampling rate that the FBD method inferred from our data, ψ = 0.01531, which implies that a species has a 31% probability of being represented in the fossil record, seems extremely high. All this cautions against attaching too much weight to our estimates of diversification and turnover.

**Table 3.** Optimised Fossilised Birth-Death parameters for the Osmundaceae. ESS = effective sample size (rounded to full numbers). For comparison, the speciation and extinction rates estimated from the unpartioned neontological data with TREEPAR (Stadler, 2011) under a model of density dependence or a constant rate model of diversification. NA = non-applicable.

| Parameter | Median value | 95% HPD interval | ESS | TREEPAR constant rate |
| --- | --- | --- | --- | --- |
| Speciation rate ($\lambda$) | 0.0299 | 0.0099–0.0549 | 10823 | 0.0314 |
| Extinction rate ($\mu$) | 0.0240 | 0.0039–0.0495 | 10890 | 0.0339<sup>b</sup> |
| Fossil recovery rate ($\psi$) | 0.0153 | 0.0065–0.0253 | 8475 | NA |
| Extant species sampling probability ($\rho$) | | = 1<sup>a</sup> | | 1 |

<sup>a</sup> In the most recent release of the FDPPDIV code that we used, $\rho$ is set to 1 by default.
<sup>b</sup> With few tips (in our case 13), TREEPAR sometimes infers negative diversification.

## CONCLUSIONS

Recent work has made clear the desirability of analysing molecular and fossil data together to jointly estimate speciation and extinction times of fossil species, branching ages of the phylogeny, and a common underlying birth-death process (Ronquist et al. 2012a, Slater et al. 2013, Silvestro et al. 2014). As shown by our trials with just 10%, 25%, and 50% of the total 36 fossils, the FBD approach is relatively insensitive to fossil sampling density (as found by Heath et al. 2014). It is, however, extremely memory craving; our data required 8 CPU processors and MCMC lengths of 20 million generations to reach stationarity after 7 hrs/run. The TE approach, at least for Osmundaceae, yielded such unexpected divergence times (and also tree topology) that we are distrustful of its results. The unexpectedly deep divergence times obtained with the FBD approach (compared to traditional node dating) caution against over-reliance on just one approach for tree calibration, but its results are more consistent with the available palaeontological record than the other trialled methods.

## ONLINE SUPPLEMENTARY MATERIAL

Online supplementary material is available for anonymous download at www.palaeogrimm.org/data/Grm14_OS.zip.

Figure. S1. The ML tree for extant Osmundaceae from the same data as used for the tree in Fig. 1 with bootstrap support at the nodes.

Figure. S2. Chronograms obtained with the FBD approach and 19 fossil rhizomes or 17 fossil fronds, and with the TE approach and 19 fossil rhizomes.

Table S1. Fossils of modern Osmundaceae used in the analyses. Table S2. List of all osmundaceous frond fossils evaluated.

Table S3. Divergence times with 95% highest posterior densities obtained from the fossilised birth-death (FBD) approach, node dating (ND) with oldest fossils, and total evidence (TE), using all 36 fossils, only the 19 fossil rhizomes, or only the 17 fossil fronds.

Table S4. Full results of the resampling analyses.

## ACKNOWLEDGEMENTS

We thank Frank Anderson, Tom Near, Tracy Heath and Daniele Silvestro for constructive criticism on the first version of the manuscript. SM acknowledges funding from the Swedish Research Council.

